# Enabling Plasmid-based Expression in *Clostridium kluyveri* using a Biparental Methylation-Conjugation System

**DOI:** 10.1101/2025.05.30.657078

**Authors:** Ethan Agena, Abiali A. Badani, Blake G. Lindner, Ian M. Gois, Andrew W. Dempster, Nigel P. Minton, Radhakrishnan Mahadevan, Christopher E. Lawson

**Affiliations:** The University of Toronto, Department of Chemical Engineering and Applied Chemistry, Toronto, ON M5T 3E5, Canada; The University of Nottingham, Synthetic Biology Research Centre, Nottingham, NG7 2RD, United Kingdom; The University of Toronto, Institute of Biomedical Engineering, Toronto, ON M5S 3E3, Canada

**Author notes:** **Corresponding Author:** Christopher E. Lawson – The University of Toronto, Department of Chemical Engineering and Applied Chemistry, Toronto, ON M5T 3E5, Canada.

**Keywords:** *Clostridium kluyveri*, methylomics, conjugation, anaerobic fermentation, microbial chain elongation, restriction-modification systems, non-model microbes

## Abstract

*Clostridium kluyveri* is a promising biocatalyst for producing medium-chain fatty acids (MCFAs) from waste-derived carbon via chain elongation. MCFAs are platform chemicals with diverse applications across agriculture, food, cosmetics, and fuels, and could support efforts towards tandem resource recovery and sustainable chemical production. However, genetic intractability has hindered efforts to engineer *C. kluyveri* for improved product yields, control over chain length and selectivity, and production of non-native oleochemicals. Here, we report a streamlined, biparental methylation-conjugation system developed for *C. kluyveri* DSM555^T^ to bypass the organism’s restriction-modification barriers and enable stable plasmid delivery. We use this system to demonstrate heterologous expression of the Fluorescence-Activated absorption-Shifting Tag (FAST), an anaerobic fluorescent reporter. This system supports advances in metabolic engineering of *C. kluyveri* and the broader adoption of genetic tools in chain elongating bacteria to expand the applications of anaerobic chain elongation in industrial biomanufacturing.

**Lay Summary:** We developed a streamlined method for genetically modifying *Clostridium kluyveri*, a promising strain for industrial anaerobic biomanufacturing, and used it to express a fluorescent protein useful for downstream applications.

**Graphical Abstract:** 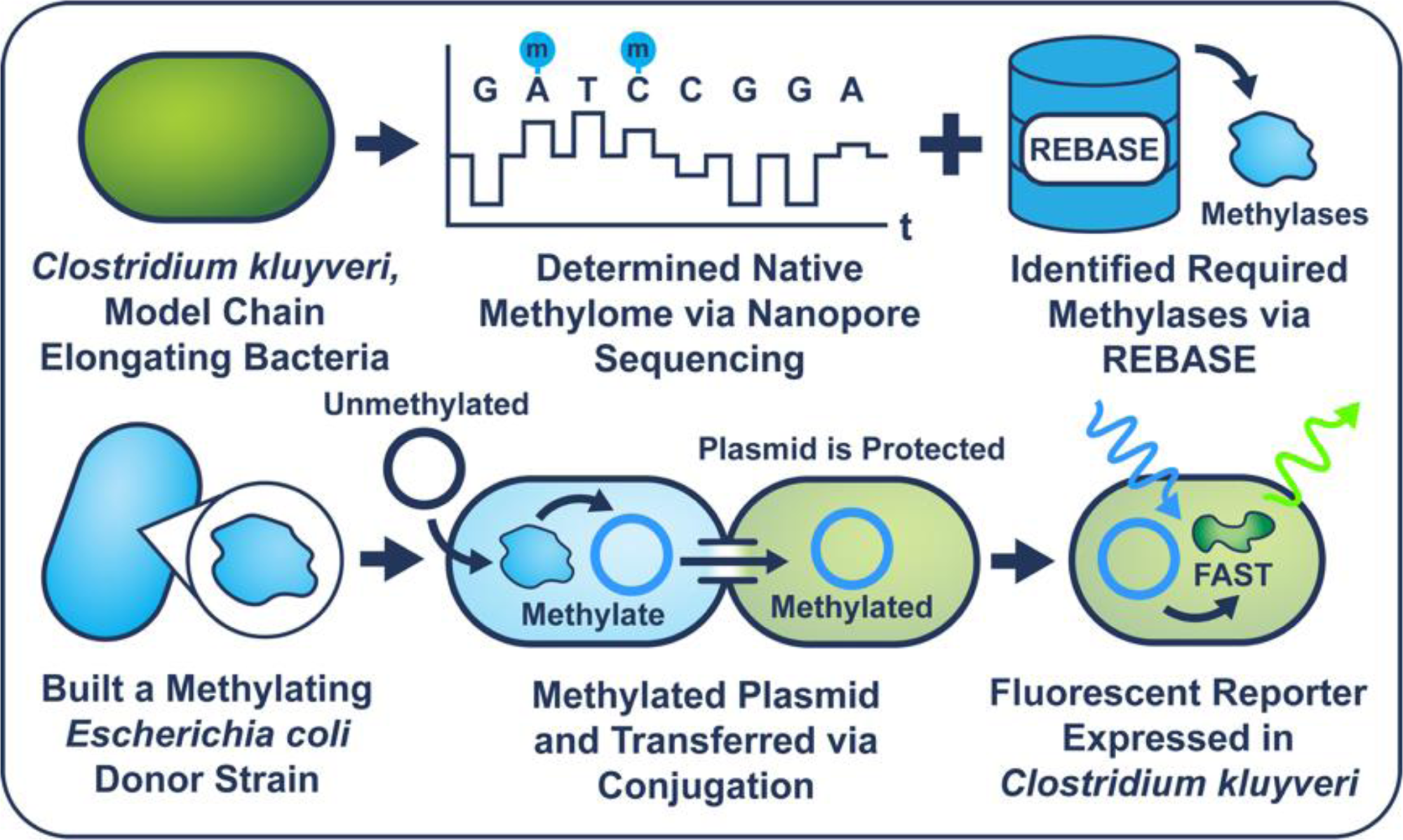

## Introduction

Anaerobic biotechnologies that transform waste carbon into high-value chemicals offer a sustainable platform for industrial biomanufacturing.^1,2^ Among these, microbial chain elongation produces C_6_–C_8_ medium chain fatty acids (MCFAs) from diverse waste streams—including agricultural residues, organic waste, and industrial syngas.^3–7^ In microbial chain elongation, chain elongating bacteria (CEB) use the reverse β-oxidation (RBO) cycle, a growth-coupled anaerobic metabolism, to produce MCFAs using waste-derived electron donors.^3,8,9^ While MCFAs have broad applications in agriculture^10^, food and cosmetic products^11–13^, and specialty chemicals and fuels^14^, current chain elongation systems face limitations in product yield, selectivity, and versatility.^11,15,16^ Improving the technology’s productivity and expanding its product range, particularly towards higher-chain MCFAs (C_9_-C_12_) and their derivatives, necessitates the development of genetic tools for CEB to manipulate native metabolic pathways and introduce heterologous functions.

Isolated in the 1940s, *Clostridium kluyveri* is the model CEB for ethanol-based chain elongation.^17^ It has been extensively studied to elucidate chain elongation physiology^3,18–20^ and applied in pure culture for the bioproduction of C_6_-C_8_ MCFAs^3,5,21^ and in microbial consortia for C_4_-C_6_ alcohols^13,22–24^. However, a lack of tools to genetically manipulate *C. kluyveri* limits deeper understanding of its metabolism and prevents metabolic engineering efforts to improve productivity and expand product range. Efforts to genetically manipulate *C. kluyveri* are impeded by its native restriction-modification (RM) systems^25^, which are bacterial defense mechanisms that protect cells from exogenous DNA (e.g., bacteriophage genomes and plasmids) by recognizing and cleaving sequences lacking host-specific methylation patterns.^26^ Type I, II, and III RM systems encode DNA methyltransferases that methylate host DNA at specific recognition sites, along with restriction endonucleases that cleave unmodified DNA; in contrast, Type IV systems encode only restriction endonucleases that target DNA with non-native methylation patterns.^27^ In effect, improperly methylated DNA is degraded by RM systems upon introduction into a host cell, which poses a critical barrier to DNA delivery for genetic manipulation.^28^

Overcoming RM systems using targeted DNA methylation is a demonstrated strategy for enabling plasmid delivery via conjugation and electro-transformation in other *Clostridium* species.^29–34^ In contrast to electro-transformation, conjugation may confer some partial evasion from RM systems by delivering single-stranded DNA, which restriction endonucleases cannot cleave.^35,36^ Concurrent work has combined these strategies and demonstrated a triparental conjugation system for *C. kluyveri*, which relies on a methylating *Escherichia coli* strain and a conjugative *E. coli* helper strain for plasmid mobilization into *C. kluyveri*.^37^ A drawback of triparental conjugation systems is the additional complexity of maintaining two separate *E. coli* strains and the reliance on two separate conjugation events, which may prolong mating times, increase mutations in the mobilized plasmids, and lead to variable transconjugant recovery.^37,38^ To complement this recent development, we report a streamlined biparental methylation-conjugation system for *C. kluyveri* DSM555^T^. Our approach uses a single *E. coli* donor strain engineered for both methylation and conjugation, following a workflow established for other *Clostridium* species.^39^ Using this system, we demonstrate plasmid transfer and functional expression of the Fluorescence-Activated absorption-Shifting Tag (FAST)^40^ anaerobic fluorescent reporter in *C. kluyveri,* an essential tool for quantifying protein expression dynamics and characterizing regulatory elements. This system opens the doors for genetic techniques to knockout/knock-in genes that could enable better control of MCFA production and access to other product classes^41^, expanding the potential applications of *C. kluyveri* beyond what is possible without metabolic engineering.

## Results

To enable exogenous plasmid delivery into *C. kluyveri* DSM555^T^, we first characterized its native RM systems via analysis of genomic methylation patterns (methylome) determined using long-read nanopore sequencing. The *C. kluyveri* methylome contains five significant motifs, which were assigned with a putative *C. kluyveri* RM system, by comparison to the entry for *C. kluyveri* on the Restriction Enzyme Database (REBASE)^42^ and all other characterized systems. This allowed us to devise a strategy to protect against the predicted RM systems, summarized in Table 1. Only motifs linked to restriction endonuclease domains required pre-methylation for effective plasmid transfer. Among the observed motifs, only the CCGG motif was associated with a Type II RM system containing a discrete restriction-endonuclease (CKL_2670) and a corresponding DNA methyltransferase (CKL_2671) amenable to pre-methylating exogenous plasmid DNA. The remaining RM systems predicted by REBASE (S1) are unlikely to pose significant barriers for plasmid transfer. Specifically, the CKL_2595 restriction endonuclease domain lacks an associated methylase gene required for its activity^43,44^ and has no unaccounted motif in the methylome, suggesting it is inactive. The other systems encode fused (CKL_2694 & CKL_2332) or multimeric (CKL_3239) methylase-restriction domains with competing activities that preclude effective restriction.

**Table 1:**
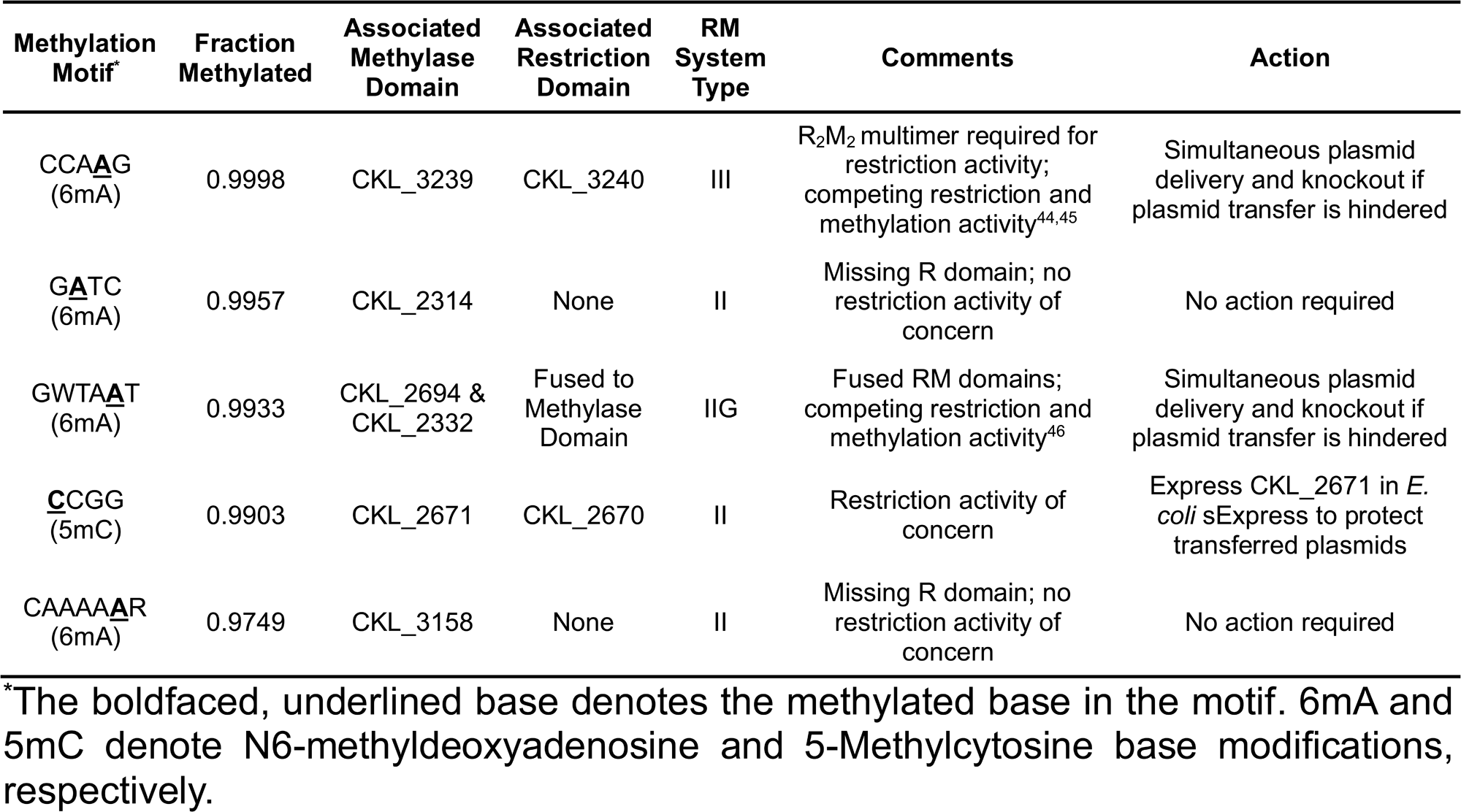
Significant methylation motifs in *C. kluyveri* DSM555^T^, their associated RM systems, and strategy for the methylation-conjugation system.

| Methylation Motif | Fraction Methylated | Associated Methylase Domain | Associated Restriction Domain | RM System Type | Comments | Action |
| --- | --- | --- | --- | --- | --- | --- |
| CCA**A**G (6mA) | 0.9998 | CKL_3239 | CKL_3240 | III | R<sub>2</sub>M<sub>2</sub> multimer required for restriction activity; competing restriction and methylation activity<sup>44,45</sup> | Simultaneous plasmid delivery and knockout if plasmid transfer is hindered |
| G**A**TC (6mA) | 0.9957 | CKL_2314 | None | II | Missing R domain; no restriction activity of concern | No action required |
| GWTA**A**T (6mA) | 0.9933 | CKL_2694 & CKL_2332 | Fused to Methylase Domain | IIG | Fused RM domains; competing restriction and methylation activity<sup>46</sup> | Simultaneous plasmid delivery and knockout if plasmid transfer is hindered |
| **C**CGG (5mC) | 0.9903 | CKL_2671 | CKL_2670 | II | Restriction activity of concern | Express CKL_2671 in *E. coli* sExpress to protect transferred plasmids |
| CAAAA**A**R (6mA) | 0.9749 | CKL_3158 | None | II | Missing R domain; no restriction activity of concern | No action required |

\*The boldfaced, underlined base denotes the methylated base in the motif. 6mA and 5mC denote N6-methyldeoxyadenosine and 5-Methylcytosine base modifications, respectively.

To simultaneously protect plasmids from the CCGG-targeting RM system in *C. kluyveri* and enable exogenous plasmid delivery, we integrated the CKL_2671 methylase into the conjugal donor strain *E. coli* sExpress, previously shown to be an effective donor strain for other *Clostridium* species.^47^ Figure 1 outlines the strain construction workflow and functional validation of the methylating conjugal donor. The CKL_2671 methylase gene was PCR-amplified from the *C. kluyveri* DSM555^T^ genome and integrated into the IncPβ conjugative plasmid R702 via λ-Red homologous recombination (S1) and transformed into *E. coli* Express to generate the methylating conjugal donor, *E. coli* sExpress CKL_2671. To assess whether *E. coli* sExpress CKL_2671 could protect plasmids from CCGG-targeting restriction endonucleases, we co-transformed it with pMTL82151, from the pMTL80000 *Clostridium* shuttle plasmid system^48^, and subjected it to restriction digestion assays. pMTL82151 carries a Gram-positive replicon from pBP1^49^ (*Clostridium botulinum*), an antibiotic resistance marker for thiamphenicol (*catP*), an IncPβ-compatible origin of transfer (*oriT)*, and 14 CCGG sites (Figure 1D). To validate *in vivo* methylation of pMTL82151 using *E. coli* sExpress CKL_2671, plasmid DNA was extracted from the strain and digested with MspI, which cleaves at CCGG sites only if the first cytosine is unmethylated.^50^ Both R702 and pMTL82151 were protected from complete digestion compared to the unmethylated control, confirming that the methylase is functional in *E. coli* (Figure 1D) and is responsible for methylating **<u>C</u>**CGG motifs in *C. kluyveri*.

**Figure 1:**
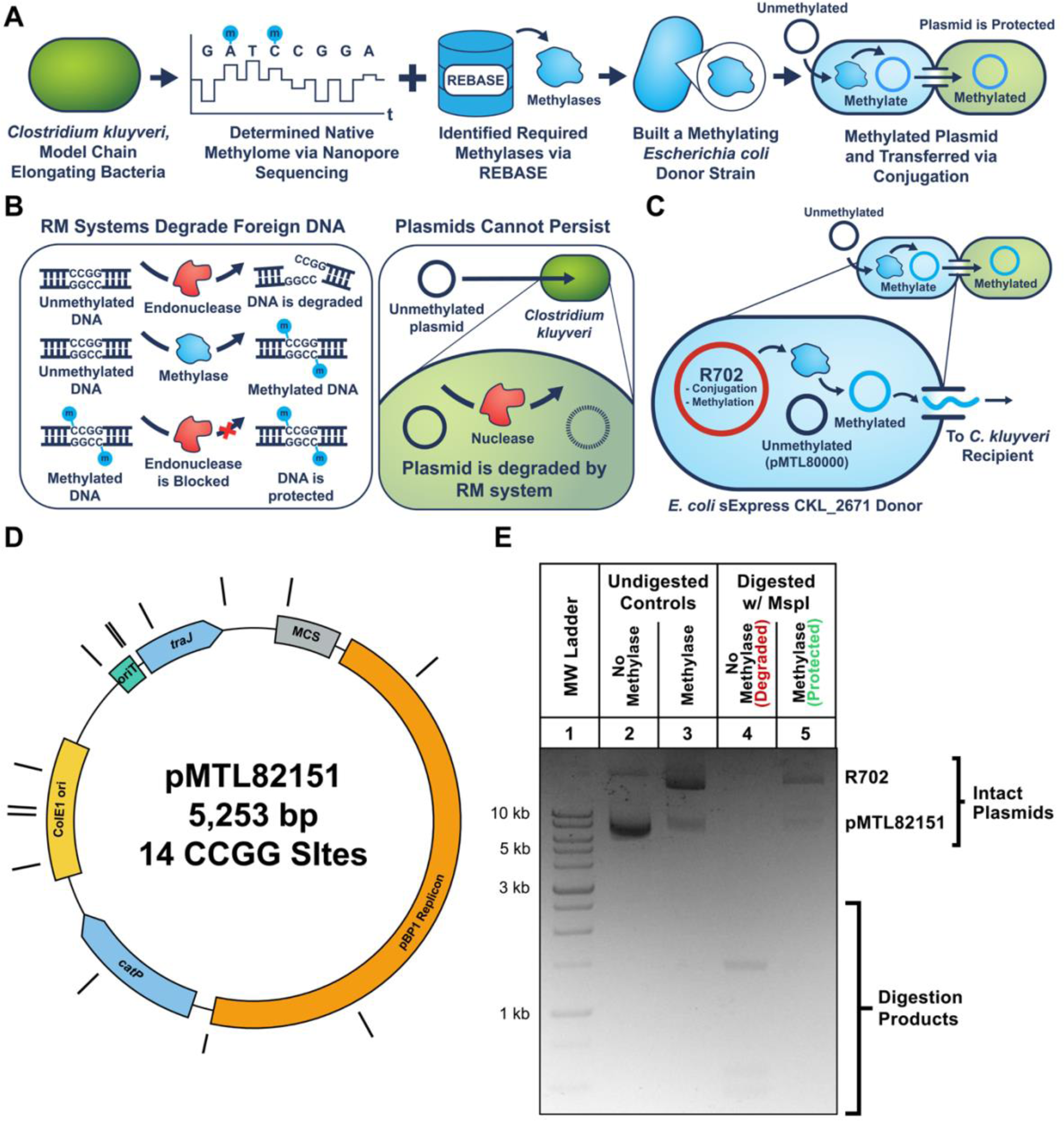
Development and functional validation of an *E. coli* methylation-conjugation strain. (A) Workflow for identifying and constructing *E. coli* donor strains expressing DNA methylases. Candidate methylases were selected based on methylome data and REBASE-annotated RM systems. (B) Conceptual overview of host RM systems, which degrade exogenous plasmid DNA lacking appropriate methylation. Plasmids propagated in standard *E. coli* are unmethylated at CCGG sites and are degraded upon entry into *C. kluyveri*. (C) Functional mechanism of the engineered *E. coli* sExpress CKL_2671 strain, which expresses the CKL_2671 methylase to methylate plasmids before conjugation. (D) Annotated plasmid map of pMTL82151, including features relevant for conjugation and selection: Multiple cloning site (MCS), pBP1 replicon, *catP*, ColE1 origin, *oriT*, and *traJ*. CCGG restriction sites targeted by MspI are highlighted as black bars. (E) MspI digestion products of plasmids extracted from *E. coli* strains with or without CKL_2671. Lane 1: Molecular weight marker (FroggaBio), Lanes 2-3: undigested controls; Lanes 4-5: plasmids digested with MspI. Intact plasmid in Lane 5 confirms functional expression of CKL_2671 *in vivo*.

Following *in vivo* methylation, we tested whether *E. coli* sExpress CKL_2671 carrying pMTL82151 could transfer thiamphenicol resistance to *C. kluyveri*. Figure 2 summarizes the results of conjugation and phenotypic characterization of the resulting *C. kluyveri* transconjugants. Using *E. coli* sExpress CKL_2671 as the donor, we successfully transferred pMTL82151 to *C. kluyveri*, isolating transconjugants under thiamphenicol selection and D-cycloserine donor counterselection (Figure 2A). Wild-type *C. kluyveri* has no intrinsic resistance to thiamphenicol, predicted using the Comprehensive Antibiotic Resistance Database^51^ and confirmed with controls. This validates the successful transfer of pMTL82151 and stable expression of the *catP* resistance gene. No colonies were obtained in the absence of methylation, demonstrating the necessity of pre-methylation for effective conjugation. Next, we used the strain to deliver protein expression plasmids and evaluated the expression of the anaerobic fluorescent reporter FAST in *C. kluyveri*. Two expression plasmids driven by the *Clostridium pasteurianum* ferredoxin promoter (P*_fdx_*) were constructed and separately co-transformed into *E. coli* sExpress CKL_2671: pMTL82153 harbouring the *lacZ α-*subunit gene (as a non-fluorescent control), and pMTL82153-FAST encoding FAST. Upon conjugation into *C. kluyveri* (Figure 2B), the identities of the *C. kluyveri* transconjugants were confirmed by 16S rRNA gene amplicon sequencing and whole plasmid sequencing. The transconjugants were cultured and incubated with the required fluorophore for FAST fluorescence, 4-hydroxy-3-methylbenzylidene-rhodanine (HMBR).^40^ Cultures expressing FAST exhibited visible yellow-green fluorescence under blue light excitation. In contrast, the wild-type and the non-fluorescent control cultures exhibited no visible fluorescence aside from red autofluorescence (Figure 2C). Quantitative fluorescence measurements further confirmed that FAST-expressing *C. kluyveri* cultures displayed significantly higher fluorescence compared to controls, with a peak emission at 542 nm (Figure 2D-E).

**Figure 2:**
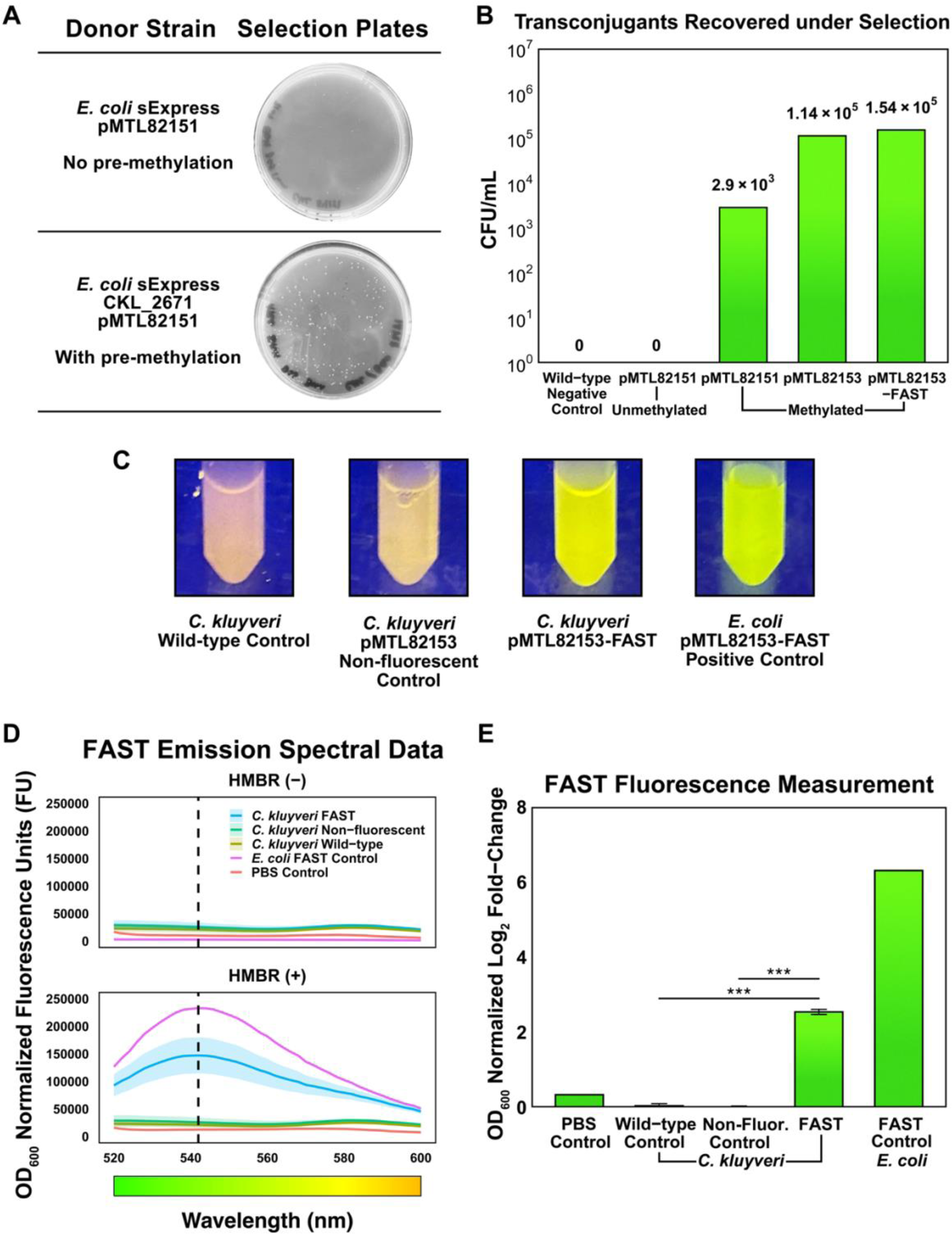
Demonstrating plasmid delivery and functional reporter expression in *C. kluyveri* DSM555^T^. (A) Representative petri plate images of *C. kluyveri* DSM555^T^ transconjugants recovered under thiamphenicol selection, indicating successful plasmid transfer. (B) Quantification of transconjugant recovery demonstrates that methylation with CKL_2671 enables efficient DNA delivery, achieving >10³ CFU/mL. (C) *C. kluyveri* cultures imaged under blue light illumination exhibit visible fluorescence from strains expressing FAST. (D) Whole cell emission spectra (Ex. 485 nm) for cultures with and without FAST expression and HMBR (shaded regions denote standard deviations). The peak wavelength for whole cells expressing FAST was measured at 542 nm. (E) Quantitative fluorescence measurements (Ex. 485 nm, Em. 542 nm) from plate reader assays show significant fluorescence in strains expressing FAST. One-way ANOVA (p = 8.42 × 10⁻⁸, α = 0.05) and Tukey’s HSD test (α = 0.05, p-values < 0.001). Plasmids: pMTL82151 (base plasmid), pMTL82153 (non-fluorescent control plasmid expressing *lacZ α-*subunit), and pMTL82153-FAST (reporter plasmid expressing FAST). Wild-type: no plasmid control.

## Discussion

This study establishes a foundational tool for the genetic engineering of the model chain elongating bacterium *C. kluyveri* DSM555^T^. By characterizing its methylome using long-read nanopore sequencing and linking key motifs to predicted RM systems, we identified and validated the activity of the CKL_2671 methylase and used it to construct a methylating conjugal donor strain, *E. coli* sExpress CKL_2671. Using this strain, we demonstrated a streamlined biparental conjugation system for *C. kluyveri*, enabling stable plasmid transfer and heterologous gene expression. Our approach achieved efficient recovery of transconjugants and exhibits several key distinctions from other systems. Unlike triparental conjugation systems^34,37,38^ that require separate *E. coli* donor and helper strains, our single-donor system simplifies the workflow and reduces the complexity of maintaining multiple strains. Additionally, we demonstrated that the pBP1 Gram-positive replicon is compatible for stable plasmid maintenance in *C. kluyveri,* expanding the plasmid toolkit for *C. kluyveri*. Importantly, we validated the functional expression of the anaerobic fluorescent reporter FAST, with engineered *C. kluyveri* strains exhibiting significantly higher fluorescence than control cultures. This marks an important step toward tunable heterologous protein expression in this organism and provides a tool for *in vivo* gene expression monitoring. To support broader adoption, the methylating donor strains developed in this study are available upon request (Associated Content).

While conjugation proved effective in our system, future efforts may benefit from developing an electroporation protocol for *C. kluyveri* to reduce the time-intensive nature of conjugation-based workflows. Although not tested here, the use of the commercially available MspI methyltransferase to pre-methylate plasmid DNA or the extraction of methylated plasmids from *E. coli* expressing CKL_2671 may circumvent RM barriers and enable electro-transformation. If successful, electroporation could offer a faster, more scalable alternative for DNA delivery into *C. kluyveri*, particularly for applications requiring iterative engineering or high-throughput library screening. Deactivating the RM systems in *C. kluyveri* via targeted knockouts could further simplify plasmid transfer into the strain, although such strategies are not applicable for engineering *C. kluyveri* within microbial consortia. Methylation-guided plasmid delivery remains critical for *in situ* engineering native microbial consortia^52^, where preserving wild-type genotypes is needed for community stability and function.

Further, the workflow for creating the methylating donor strain can be extended to other CEB once methylomes for these organisms are determined. Future efforts will focus on expanding the toolkit for *C. kluyveri*, including tunable and inducible expression systems, using emerging fluorescent reporters with different emission wavelengths (e.g., evolved flavin-based fluorescent proteins (FbFPs)^53^ and anaerobic, blue fluorescent proteins^54^), which will be critical for high-throughput screening, multi-gene expression tracking, and protein-protein interaction techniques like Fluorescence resonance energy transfer (FRET)^55^ which has been inaccessible in anaerobic conditions.

Developing programmable genome editing tools for *C. kluyveri* is another important next step. Potential strategies include adapting heterologous systems for *Clostridium,* such as RiboCas^56^, leveraging *C. kluyveri*’s endogenous CRISPR-Cas system for native interference or editing capabilities^57^, and applying transposon mutagenesis libraries for high-throughput genotype-phenotype mapping.^58^ Establishing robust genome editing workflows will deepen our mechanistic understanding of chain elongation and enable metabolic engineering of *C. kluyveri*. Such tools could facilitate targeted improvements in chain elongation yield, chain length selectivity, product tolerance, and expand the product spectrum of *C.* kluyveri to other oleochemicals, increasing its utility as a chassis for sustainable industrial biomanufacturing.

## Methods

### Bacterial Strains and Culture Conditions

*E. coli* strains were grown aerobically at 37 °C in LB medium or plates with appropriate antibiotics for each strain (S1). *C. kluyveri* DSM555^T^ was cultured anaerobically in Hungate tubes containing ATCC 1120 liquid medium at 37 °C and plated on Reinforced Clostridial Medium (RCM) agar supplemented with 100 µL of ethanol spread-plated before inoculation in an anaerobic chamber (COY, N_2_/CO_2_/H_2_ atmosphere) and incubated at 37 °C in anaerobic containers (BD GasPak EZ Container Systems).

### *C. kluyveri* Methylome and RM Systems Analysis

Genomic DNA from *C. kluyveri* DSM555^T^ was extracted using the Zymo Quick-DNA HMW MagBead Kit with modifications: bead beating (0.1 mm glass beads) and mutanolysin (50 U) were added to improve lysis. DNA was further purified with TotalPure NGS Beads (0.7× ratio) to remove low-molecular-weight fragments. Sequencing libraries were prepared with the ONT Rapid Barcoding Kit V14 and run on a MinION device (R10.4.1). Basecalling was done with dorado (v0.9.5), and an assembly generated using Flye^59^ (v2.9.5), assessed with CheckM2^60^ (v1.0.2), and taxonomically validated with GTDBtk^61^ (v2.4.1). 6mA and 5mC modifications were detected via dorado’s methylation model and mapped with minimap2^62^ (-ax map-ont). Genome-wide patterns and high-confidence motifs were analyzed using modkit^63^ (v0.4.4). Motifs associated with a putative *C. kluyveri* RM system based on REBASE. For RM systems lacking known recognition motifs, we predicted likely targets based on homology to characterized RM systems in REBASE. More details can be found in the Supporting Information (S1).

### Construction of *Clostridium* Expression Plasmids

*Clostridium* shuttle plasmids for *E. coli* and *C. kluyveri* were assembled using modular parts from the pMTL80000 plasmid series from the Synthetic Biology Research Centre, University of Nottingham (SBRC). Restriction-digested or PCR-amplified fragments were assembled with NEBuilder HiFi Assembly and transformed into *E. coli* NEB5a (NEB). Final plasmids were sequence-verified by long-read sequencing (Plasmidsaurus).

### Construction of the Methylating Conjugal Donor Strain

The *E. coli* sExpress CKL_2671 methylating conjugal donor was constructed based on a novel workflow leveraged for other *Clostridium* species from collaborators at the SBRC. The workflow included amplifying the *C. kluyveri* CKL_2671 methylase gene from its genome and cloning it into the arabinose-inducible RBV-1 vector via NEBuilder HiFi assembly and transformed into *E. coli* TOP10. The CKL_2671 expression cassette in RBV-1 includes homology arms for integration into the R702 plasmid. The CKL_2671 expression cassette was PCR-amplified and integrated into R702 via λ-Red recombineering in *E. coli* RE-InterStellar (harbouring R702 and pKD46cm). Recombinants were selected on apramycin and kanamycin, verified by PCR across the insertion site, and cured of pKD46cm. The recombinant R702 (containing CKL_2671 expression cassette) was extracted and transformed into *E. coli* Express (NEB) to obtain the final methylating conjugal donor, *E. coli* sExpress CKL_2671. *E. coli* sExpress CKL_2671 was co-transformed with the assembled *Clostridium* shuttle plasmids via heat-shock transformation. Functional methylation of these conjugal donors was confirmed by protection against MspI digestion (methylation-sensitive).

### Anaerobic conjugation and validation of transconjugants

*E. coli* sExpress CKL_2671 conjugal donors strains carrying the *Clostridium* shuttle plasmids were mated with *C. kluyveri* under anaerobic conditions. *E. coli* and *C. kluyveri* were mixed at a donor:recipient ratio of 1:1, spotted on RCM agar supplemented with ethanol, and incubated for 48 h. Total biomass of the conjugal mixture was harvested and plated at several serial dilutions on RCM agar supplemented with ethanol along with thiamphenicol [7.5 μg/mL] to select for *C. kluyveri* transconjugants and D-cycloserine [250 μg/mL] to select against the *E. coli* donor strain. Transconjugants colonies were screened after 5 d by colony morphology, inoculated into Hungate tubes containing ATCC1120 supplemented with thiamphenicol and D-cycloserine at the same concentrations above and incubated at 37 °C for 3 d. Transconjugant cultures were passaged once, and nucleic acids were extracted for 16S rRNA gene sequencing to confirm strain identity, and long-read sequencing to confirm plasmid identity.

### FAST fluorescence measurement

*E. coli* and *C. kluyveri* cultures were washed with PBS and resuspended to OD_600_ = 1.0 in PBS ± 20 µM 4-hydroxy-3-methylbenzylidene-rhodanine (HMBR, stock prepared in DMSO). Fluorescence was measured in flat-bottom, black 96-well plates (Greiner, 200 µL/well) using a plate reader and normalized to the measured OD_600_ (Tecan Infinite M Plex, Ex: 485 nm, Em: 542 nm).

## Associated Content

The Supporting Information is available free of charge at [insert publication link]. Supporting Information: Additional experimental details, materials, and methods (PDF). The conjugal donor strains developed as part of this work are available upon request at cost.

## Author Information

Complete contact information is available at: [insert publication link]

### Author Contributions

E.A. conceived, designed, and performed all experiments, analyzed the data, and drafted the manuscript. A.A.B., B.G.L., and I.M.G. assisted with methylome analysis and interpretation. A.W.D. provided training and guidance for methylation strain construction and recombineering techniques. N.P.M., R.M., and C.E.L. supervised the project and contributed to experimental design, data interpretation, and manuscript review. All authors reviewed and approved the final manuscript.

### Notes

A.W.D. is an employee of Forge Genetics Inc., which offers the methylating *E. coli* strains described in this work at cost for academic and non-profit research. All other authors declare no competing financial interests.

## Acknowledgements

The authors thank Dr. Adam Guss (Oak Ridge National Laboratory, University of Tennessee) for his early guidance on methylome analysis strategy and interpretation. We also acknowledge Robinson Meng for his assistance with routine lab work and strain maintenance during early project stages. This work was supported by the Natural Sciences and Engineering Research Council of Canada (ALLRP 580897-22, RGPIN-2021-02684) awarded to C.E.L.

## Supporting Information (S1)

### Restriction-Modification Systems Analysis for *Clostridium kluyveri*

The REBASE Database (New England Biolabs) was used to both bioinformatically predict the presence of restriction modification systems (types I-IV) and the highest probability recognition motif of each annotated methyltransferase and endonuclease within *C. kluyveri*. Predicted enzymes were automatically assigned locus tags and sorted based on RM type, methyltransferase (M) or endonuclease activity (R), and their homology predicted recognition motif. Independently collected methylome data was further used to narrow down motifs of interest by identifying discrete genomic motifs that had a high incidence of methylation (5mC, 4mC, and 6mA).

Using both data sets, annotated REBASE motifs for *C. kluyveri* were first cross referenced with the genomic motifs and methylation type when available. For all remaining enzyme sets the motif and methylation data of closest annotated relative was used. From this comparison six potential candidates were identified with matching motif data, indicating these systems were actively being express. Of these, only four targets (CKL_2332, CKL2670/2671, CKL_2694, and CKL_3239/3240) had an annotated endonuclease that is required for DNA cleavage and degradation, indicating that they are likely barriers to conjugation and thus targets for evasion or removal. With a list of four targets, the RM type between each system was then compared to assign a rank to each system based on both likelihood of plasmid cleavage and ease of targeting.

CKL 3239/3240 was the only annotated type III system, which requires the formation of a heteromeric protein complex that facilitates both cleavage and modification of DNA motifs. Due to both the stability and dual function of the complex, and the lack of suitable commercial isozymes, isolation of methylation activity is not always possible. Thus, these systems are challenging to evade through DNA pre-methylation and require RNA interference strategies or gene knockouts to remove. Similarly, CKL_2332 and CKL_2694 are type IIG systems, consisting of a single protein with a fused restriction and modification domain, making the isolation impossible due to the significant overlap of the domains. CKL2670/2671 however, is a canonical type II system, consisting of a separated methylase and endonuclease pair with isolated modification and restriction activity. This enabled the selective pre-methylation of plasmid DNA using CKL_2671 to evade the restriction of CKL_2670 and confer a layer of epigenetic protection to plasmids pre-conjugation.

**Table S1:**
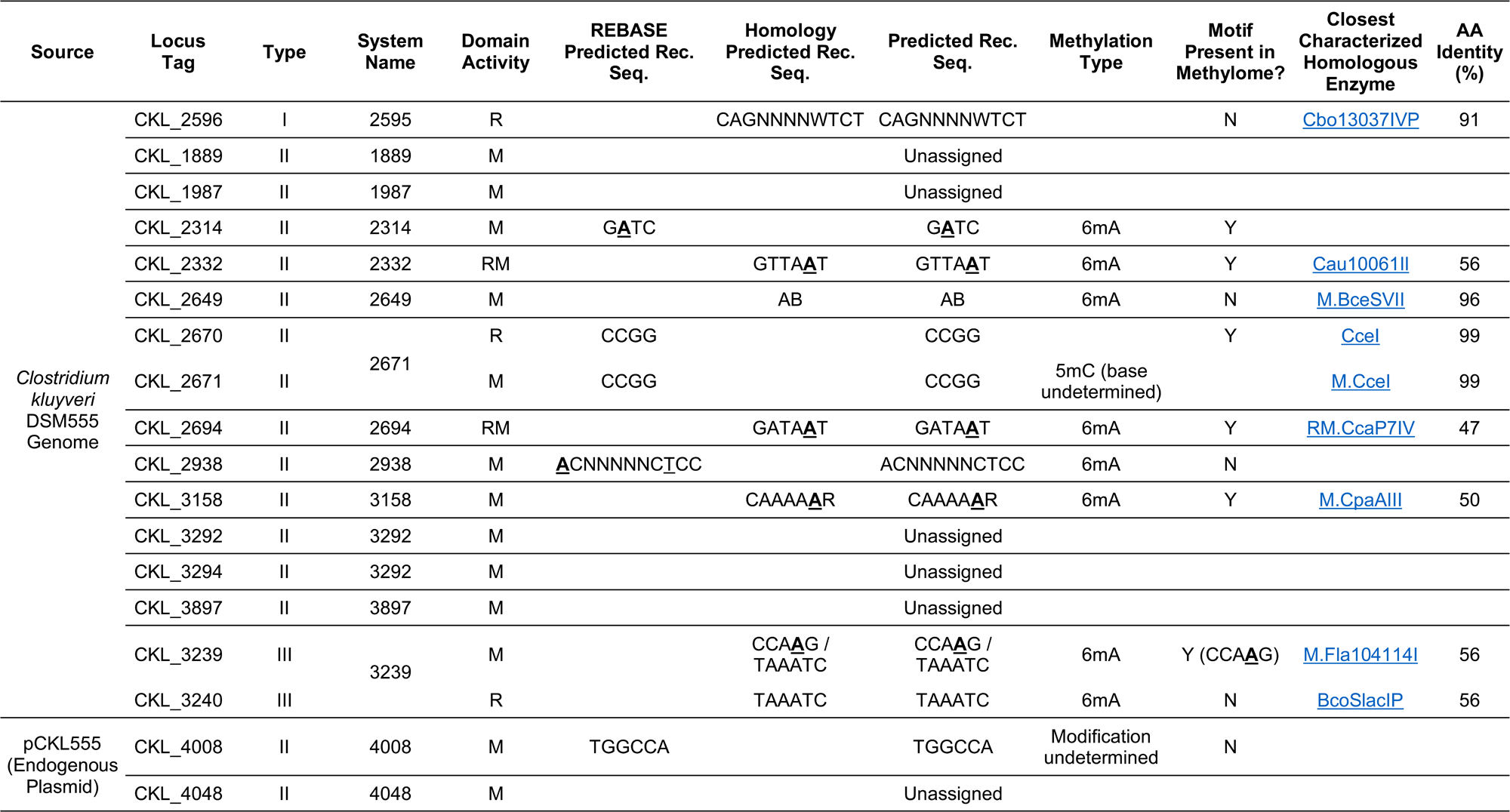
REBASE Predicted RM systems for *C. kluyveri* and recognition sequence assignment.

### Methylome Analysis of *Clostridium kluyveri*

Genomic DNA was extracted from *Clostridium kluyveri* DSM 555 cultures harvested at late exponential phase using the **Zymo Quick-DNA HMW MagBead Kit** (Zymo Research), following the manufacturer’s protocol optimized for high-molecular-weight DNA suitable for long-read sequencing. Two modifications to the recommended protocol were made to optimize genomic DNA yield and purity. First, cell pellets were subjected to bead beating with sterile, acid-washed 0.1mm glass beads. Second, a total of 50U mutanolysin (Sigma-Aldrich, M9901) was added to the enzymatic lysis buffer. Additionally, the resulting high molecular weight gDNA was furthered purified via TotalPure NGS Beads (Omega Bio-tek) using a 0.7X ratio of magnetic beads to sample volume to deplete low molecular weight DNA fragments before library preparation.

Purified gDNA was then prepared for sequencing on a minION device (Oxford Nanopore Technologies) using Rapid Barcoding Kit V14 (SQK-RBK114) library preparation followed by loading and sequencing on a minION flow cell (R10.4.1) following manufacturer’s protocols. The resulting raw data was basecalled with dorado (v0.9.5) using the relevant high-accuracy model with 400bps translocation speed (model v5.0.0). Reference genome assembly was performed with flye (v2.9.5) using default settings. Thorough quality assessment of the resulting assembly was completed with CheckM2 (v1.0.2) and confirmation of taxonomic assignment to *C. kluyveri* DSM 555 was verified with GTDBtk (v2.4.1) against GTDB release 226.

Methylation of adenine (6mA) and cytosine (4mC and 5mC) bases were sensed with dorado using appropriate modification-calling extensions of the canonical model mentioned above (dna_r10.4.1_e8.2_400bps_hac). The resulting methylated sequences were mapped to the reference genome assembly produced herein with minimap2 (-ax map-ont; v2.28). Genome-wide methylation patterns were then analyzed using modkit (v0.4.4); the proportion of mapped reads carrying modifications at each position was summarized with modkit pileup (--filter-threshold 0.7) and motifs with high proportions of modification were deduced by modkit motif search. Finally, the log2 odds of the reported high confidence motifs were calculated with modkit motif evaluate.

**Table S3:**
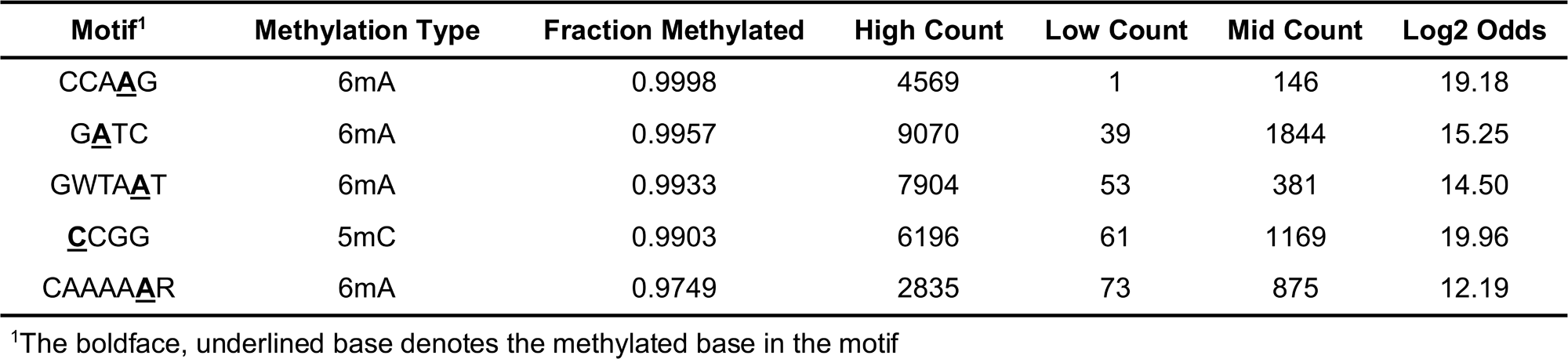
Significant methylation motifs in *C. kluyveri*.

To associate detected methylation motifs with putative restriction-modification (RM) systems, motif data were cross-referenced with REBASE, enabling annotation of *C. kluyveri* methyltransferases and endonucleases based on homology to previously characterized systems.

### Integration of CKL_2671 into R702

The workflow conducted has been recently demonstrated in other *Clostridium* species but had not been applied to *Clostridium kluyveri.* To construct the methylating conjugal donor strain, the DNA methyltransferase (methylase) gene of interest (CKL_2671) was amplified by PCR from the *C. kluyveri* genome, including the native ribosome binding site and appropriate 5’ overhangs for HiFi DNA assembly. The CKL_2671 gene was then assembled into the *E. coli*-based recombineering vector RBV-1, which contains an arabinose-inducible *pBAD* promoter controlling the methylase, a ColE1 origin of replication for plasmid maintenance in *E. coli*, homology arms for R702, and dual antibiotic markers (erythromycin and apramycin resistance).

RBV-1 was linearized by digestion with EcoRI and BsaI (NEB), and the gel-purified PCR products were inserted into the linearized vector using NEBuilder HiFi DNA assembly. The assembled plasmid was transformed into *E. coli* TOP10 cells, as other strains may harbor active Type IV restriction systems that are incompatible with methylase expression. Transformants were screened by colony PCR and confirmed by sequencing. Functional validation of methylase activity was performed using restriction digest assays with isoschizomers for the CCGG recognition sequence (MspI and HpaII, NEB) corresponding to the predicted methylation motif of CKL_2671.

Once methylation activity was confirmed, the assembled methylase integration cassette—including homology arms for R702 flanking the insertion site—were amplified and inserted into the *E. coli* R702 conjugal plasmid using lambda Red recombineering. This step employed an electro-transformation of the CKL_2671 integration cassette PCR product into *E. coli* RE-InterStellar, which harbours the R702 plasmid and pKD46cm, which carries the genes for the lambda Red system. At this stage, culturing was performed at 30°C as pKD46cm contains a temperature sensitive replicon. Recombinants were selected on LB agar containing apramycin and kanamycin, pKD46cm was cured with subsequent culturing at 37°C, and correct insertion into R702 was verified by PCR spanning the homology arm junctions and by confirming the loss of spectinomycin resistance, which serves as a negative selection marker.

Following this, the recombinant R702-CKL_2671 plasmid was extracted from *E. coli* InterStellar and electro-transformed into E. coli Express to yield the base methylating strain, E. coli sExpress CKL_2671. To create the conjugal donors with mobilizable plasmids for *Clostridium, E. coli* sExpress CKL_2671 was transformed via heat-shock with plasmids from the pMTL80000 *Clostridium* shuttle plasmid series. Again, functional validation of methylase activity was performed using restriction digest assays on plasmid extracts from the final conjugal donors. The system demonstrated here supports foundational advancements for the genetic manipulation of *C. kluyveri* DSM 555^T^ with potential for translation to *C. kluyveri* NBRC 12016, which has similar RM systems based on REBASE predictions.

**Figure S1:**
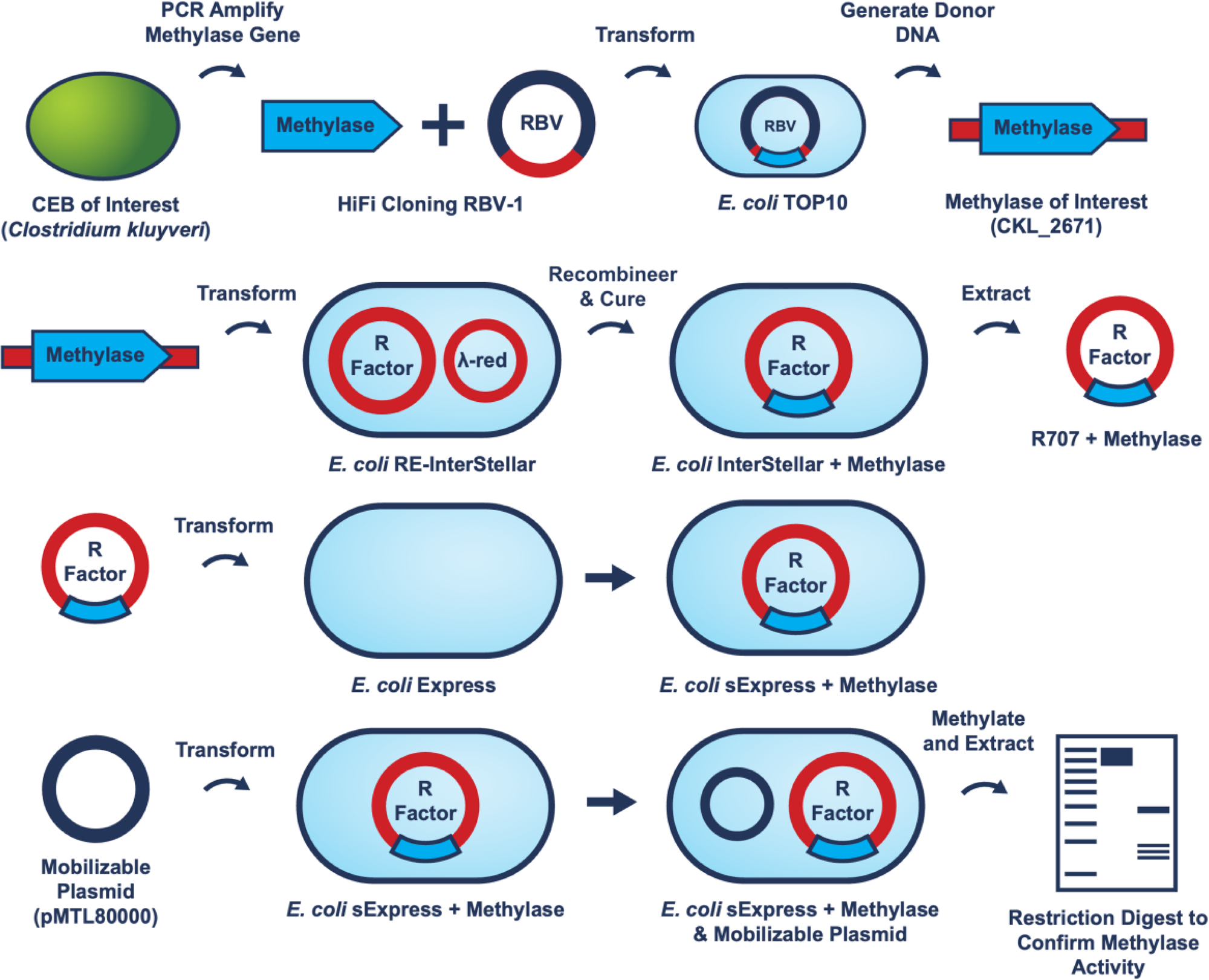
Schematic of the molecular cloning workflow for constructing the methylating conjugal donor strain, *E. coli* sExpress CKL_2671

### Bacterial Strains, Media, and Growth Conditions

**Table S2:**
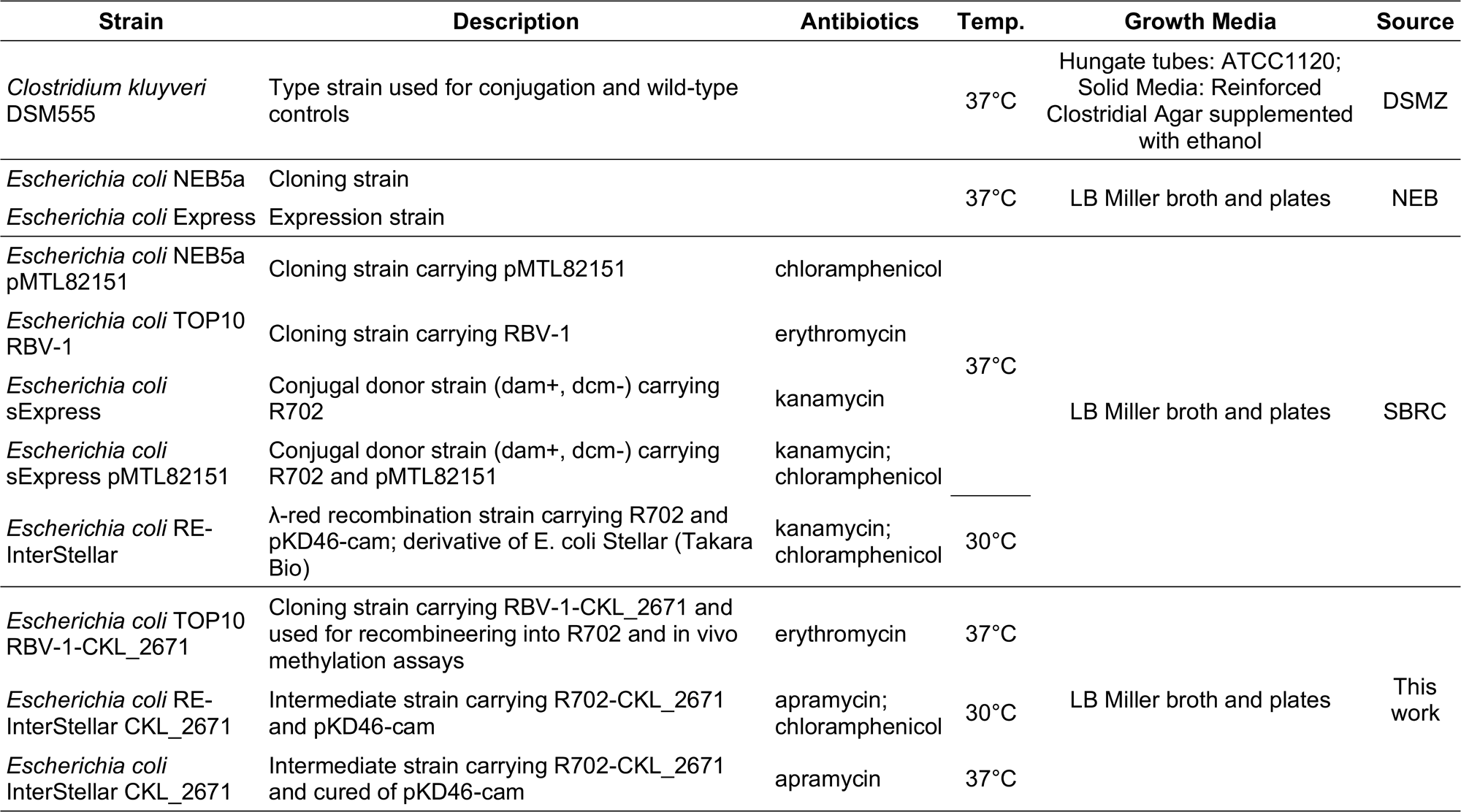
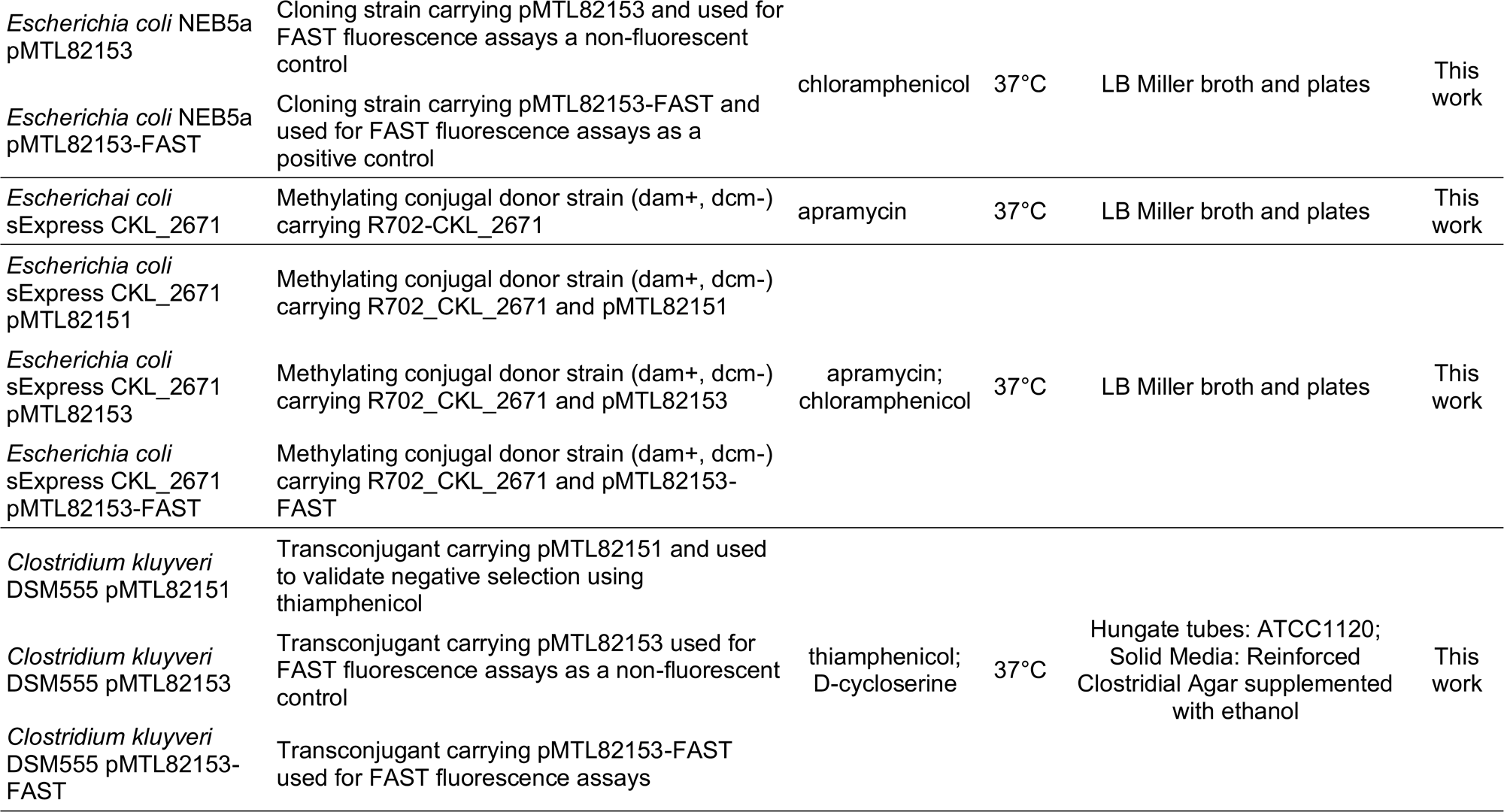
A list of bacterial strains and their growth conditions used for this study.

### Primers

**Table S4:**
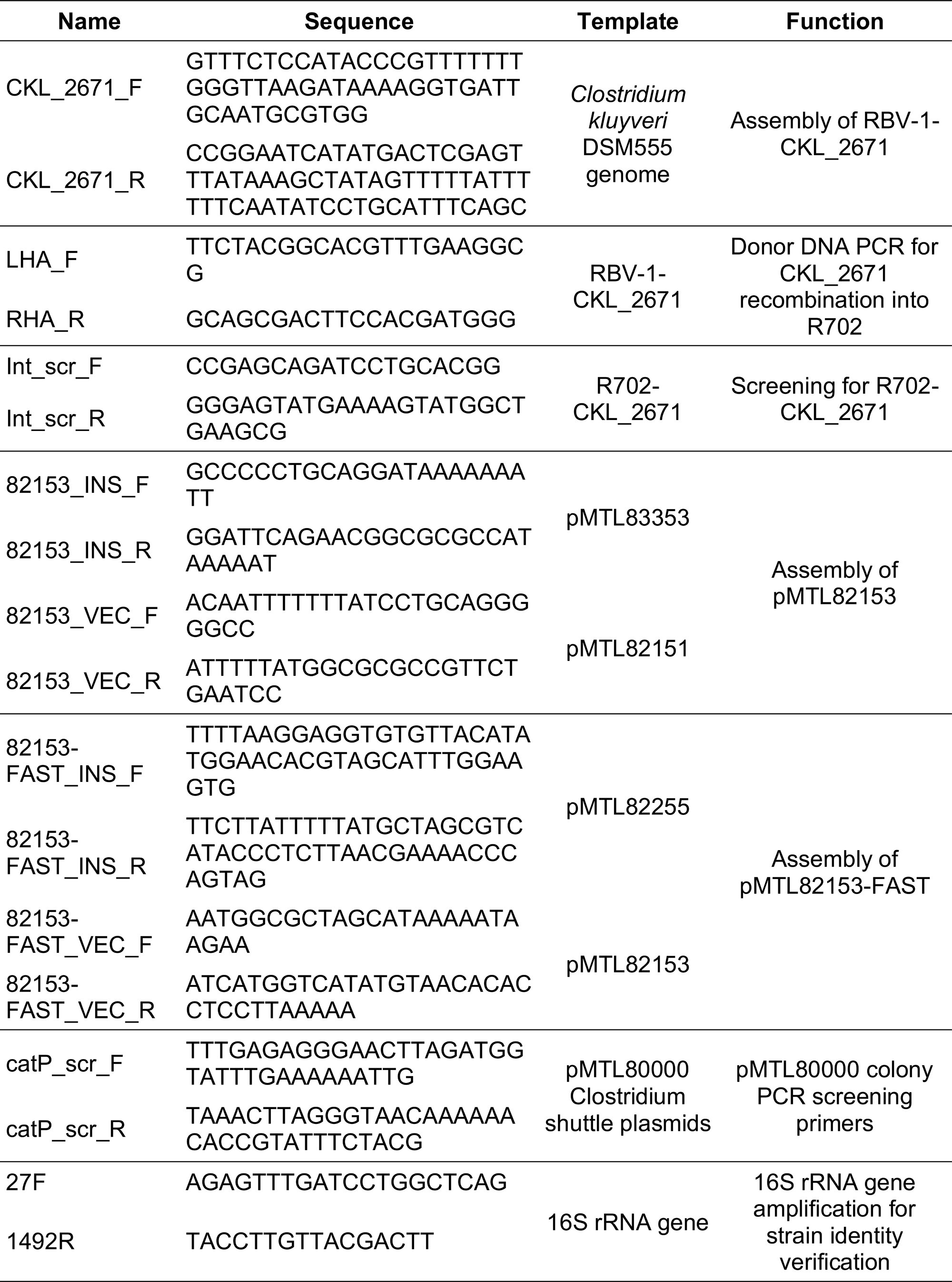
A list of primers used in this study.

### Plasmids

**Table S5:**
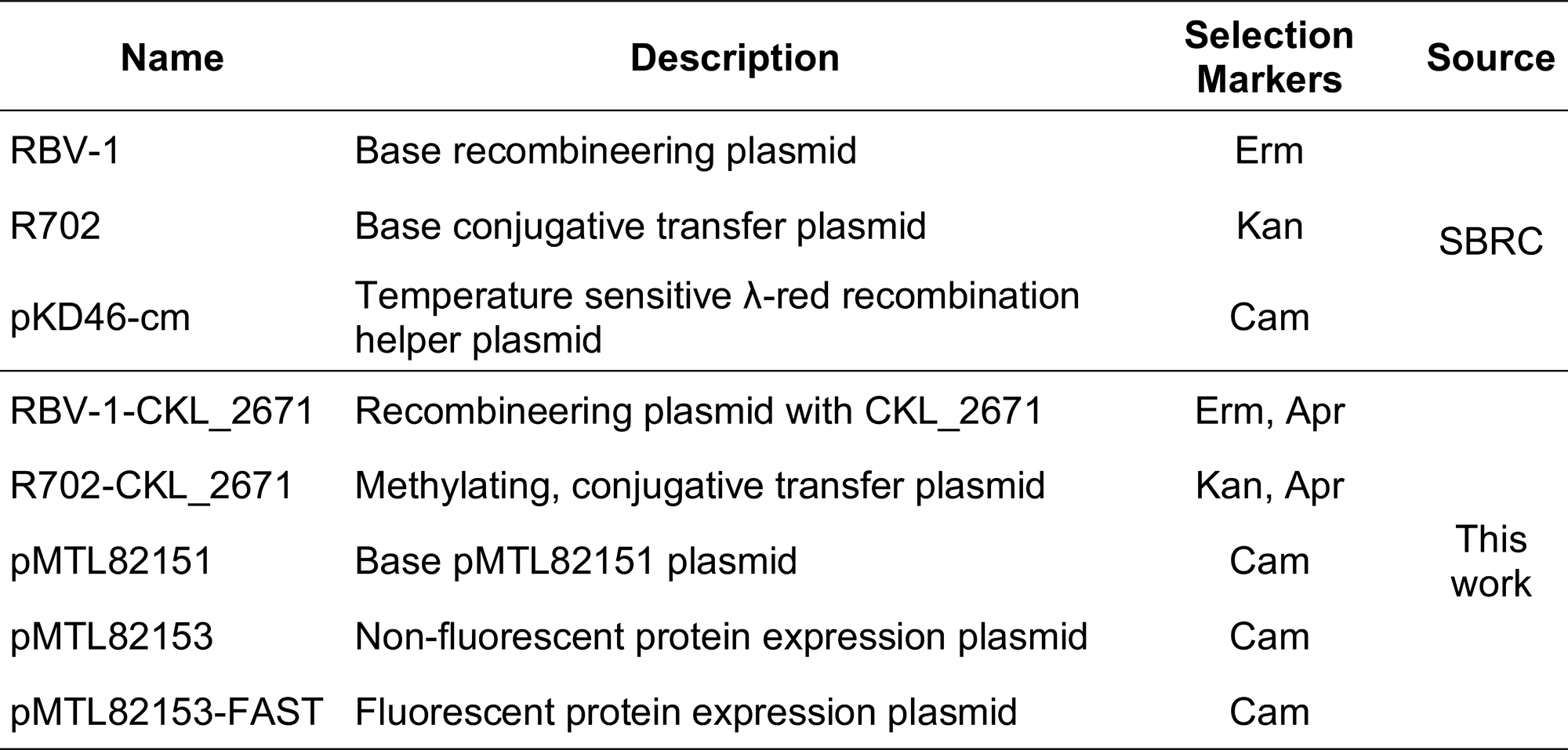
A list of plasmids used in this study.

**Figure S2:**
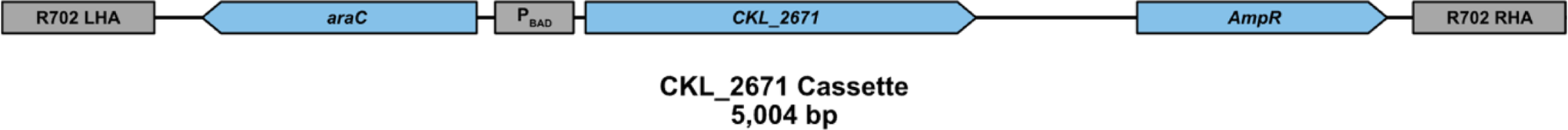
A schematic of the CKL_2671 Expression Cassette integrated into the R702 plasmid. R702 LHA: Left-hand homology arm (LHA) for the R702. araC: coding sequence for the P_BAD_ repressor protein. P_BAD_: arabinose-inducible promoter controlling CKL_2671 expression. AmpR: resistance marker for apramycin negative selection. CKL_2671: a 5mC DNA methyltransferase from the *C. kluyveri* genome, methylating **<u>C</u>**CGG sites at the first cytosine. R702 RHA: right-hand homology arm (RHA) for the R702.

**Figure S3:**
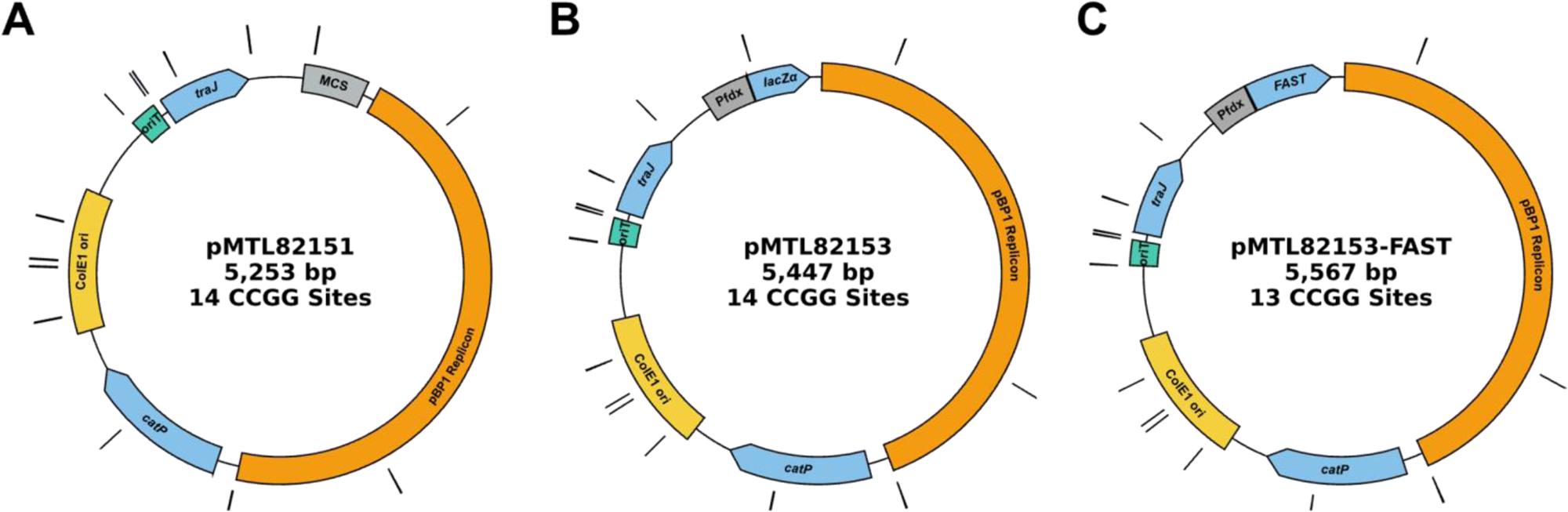
Plasmid maps of the pMTL80000 plasmids used in this study with annotated CCGG sites. pBP1 Replicon: Gram-positive compatible replicon from pBP1. *catP*: an antibiotic resistance marker for thiamphenicol. ColE1 ori: Gram-negative compatible origin of replication. *oriT:* an IncPβ-compatible origin of transfer. *traJ*: a coding sequence for the transcriptional activator for conjugative functions of R702. The plasmids differ at the insert region: (A) pMTL82151 contains a multiple-cloning site (MCS). (B) pMTL82153 contains an expression cassette for the lacZ α-subunit controlled by P_fdx_. (C) pMTL82152-FAST contains an expression cassette for FAST controlled by P_fdx._

### Characterization of the CKL_2671 DNA Methyltransferase Activity

Prior to constructing the methylation strain, we confirmed the *in vivo* activity and modification specificity of CKL_2671 in *E. coli*. CKL_2671 was cloned into the intermediate vector RBV-1 using HiFi Assembly. Expression of CKL_2671 in *E. coli* RBV-1 was induced using 10 mM arabinose, and plasmid DNA was extracted and subjected to MspI and HpaII digestion.

RBV-1-CKL_2671 was extracted and subjected to restriction digestion with MspI and HpaII, which both recognize the CCGG sequence targeted by the CKL_2671, methylase. These enzymes differ in their sensitivity to cytosine methylation: MspI is blocked by methylation of the external cytosine (**<u>C</u>**CGG), while HpaII is blocked by methylation of the internal cytosine (C**<u>C</u>**GG).

RBV-1-CKL_2671 was protected from cleavage by both enzymes, confirming *in vivo* activity of CKL_2671 and confirming methylation at the external cytosine (**<u>C</u>**CGG). The observed protection confirmed the role of CKL_2671 in the *C. kluyveri* methylome and RM system and validated it as a viable candidate for methylation strain construction.

**Figure S4:**
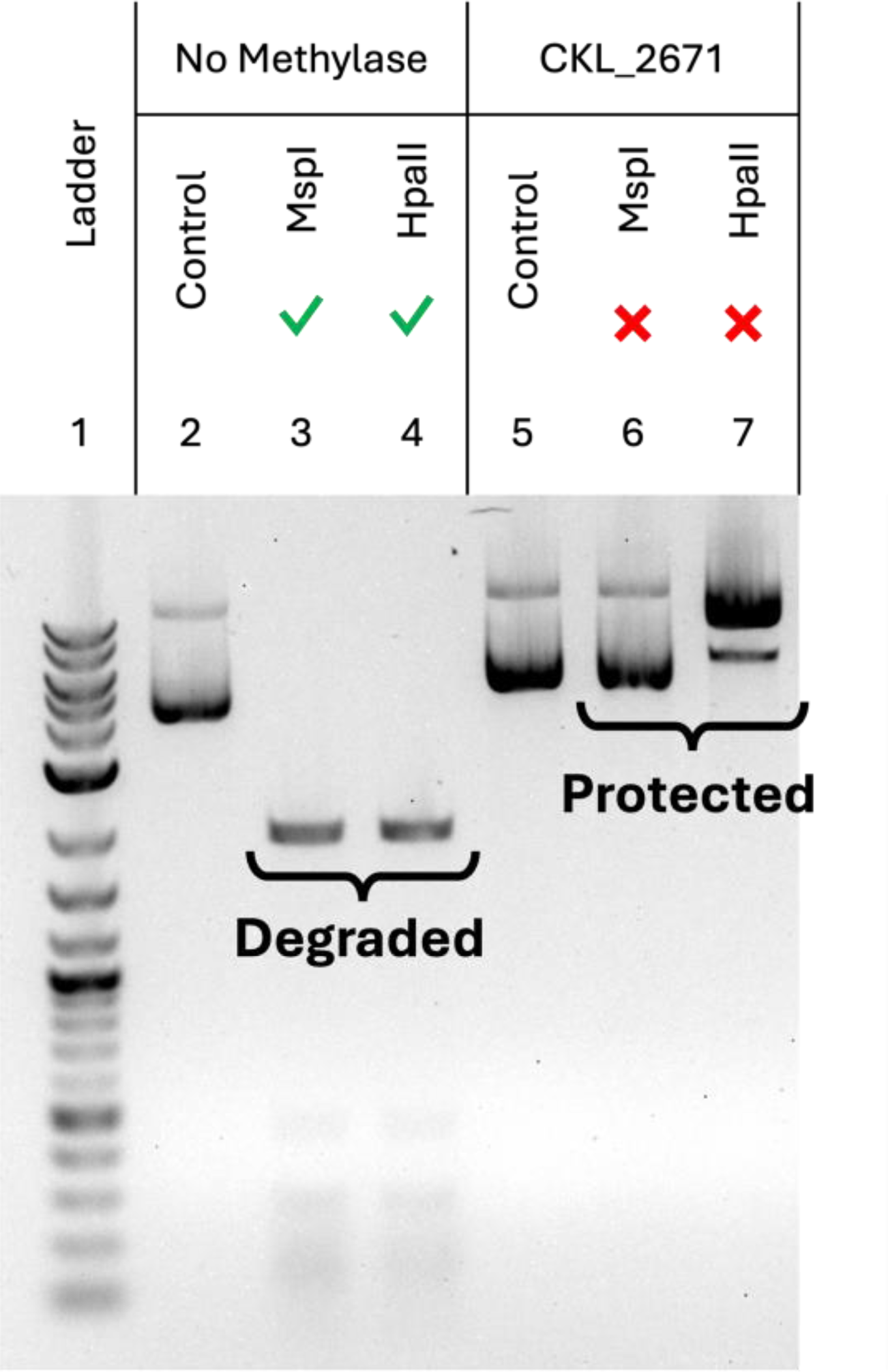
**3**The methylation sensitivity of the commercial restriction enzymes used in the digest. MspI and HpaII both recognize the sequence CCGG, but their activity is both blocked by methylation of the external cytosine. DNA gel electrophoresis image of plasmid digests (1% agarose). Lanes 2 and 5 are undigested controls. Lanes 3-4 and 6-7 are plasmids digested with MspI and HpaII with (RBV-1-CKL_2671) or without the methylase (RBV-1).

