## Supporting Information S1 for "Enabling Plasmid-based Expression in *Clostridium kluyveri* using a Biparental Methylation-Conjugation System"

Table S1: REBASE Predicted RM systems for *C. kluyveri* and recognition sequence assignment

| Source | Locus Tag | Type | System Name | Domain Activity | REBASE Predicted Rec. Seq. | Homology Predicted Rec. Seq. | Predicted Rec. Seq. | Methylation Type | Motif Present in Methylome? | Closest Characterized Homologous Enzyme | AA Identity (%) |
|---|---|---|---|---|---|---|---|---|---|---|---|
| *Clostridium kluyveri* DSM555 Genome | CKL_2596 | I | 2595 | R | | CAGNNNNWTCT | CAGNNNNWTCT | | N | Cbo13037IVP | 91 |
| | CKL_1889 | II | 1889 | M | | | Unassigned | | | | |
| | CKL_1987 | II | 1987 | M | | | Unassigned | | | | |
| | CKL_2314 | II | 2314 | M | G**A**TC | | G**A**TC | 6mA | Y | | |
| | CKL_2332 | II | 2332 | RM | | GTTA**A**T | GTTA**A**T | 6mA | Y | Cau10061II | 56 |
| | CKL_2649 | II | 2649 | M | | AB | AB | 6mA | N | M.BceSVII | 96 |
| | CKL_2670 | II | 2671 | R | CCGG | | CCGG | | Y | CceI | 99 |
| | CKL_2671 | II | | M | CCGG | | CCGG | 5mC (base undetermined) | | M.CceI | 99 |
| | CKL_2694 | II | 2694 | RM | | GATA**A**T | GATA**A**T | 6mA | Y | RM.CcaP7IV | 47 |
| | CKL_2938 | II | 2938 | M | **A**CNNNNNCTCC | | ACNNNNNCTCC | 6mA | N | | |
| | CKL_3158 | II | 3158 | M | | CAAAA**A**R | CAAAA**A**R | 6mA | Y | M.CpaAIII | 50 |
| | CKL_3292 | II | 3292 | M | | | Unassigned | | | | |
| | CKL_3294 | II | 3292 | M | | | Unassigned | | | | |
| | CKL_3897 | II | 3897 | M | | | Unassigned | | | | |
| | CKL_3239 | III | 3239 | M | | CCA**A**G / TAAATC | CCA**A**G / TAAATC | 6mA | Y (CCA**A**G) | M.Fla104114I | 56 |
| | CKL_3240 | III | | R | | TAAATC | TAAATC | 6mA | N | BcoSlacIP | 56 |
| pCKL555 (Endogenous Plasmid) | CKL_4008 | II | 4008 | M | TGGCCA | | TGGCCA | Modification undetermined | N | | |
| | CKL_4048 | II | 4048 | M | | | Unassigned | | | | |

Table S3: Significant methylation motifs in *C. kluyveri*.

| Motif[1] | Methylation Type | Fraction Methylated | High Count | Low Count | Mid Count | Log2 Odds |
|---|---|---|---|---|---|---|
| CCA**A**G | 6mA | 0.9998 | 4569 | 1 | 146 | 19.18 |
| G**A**TC | 6mA | 0.9957 | 9070 | 39 | 1844 | 15.25 |
| GWTA**A**T | 6mA | 0.9933 | 7904 | 53 | 381 | 14.50 |
| **C**CGG | 5mC | 0.9903 | 6196 | 61 | 1169 | 19.96 |
| CAAAA**A**R | 6mA | 0.9749 | 2835 | 73 | 875 | 12.19 |

[1]The boldface, underlined base denotes the methylated base in the motif

To associate detected methylation motifs with putative restriction-modification (RM) systems, motif data were cross-referenced with REBASE, enabling annotation of *C. kluyveri* methyltransferases and endonucleases based on homology to previously characterized systems.

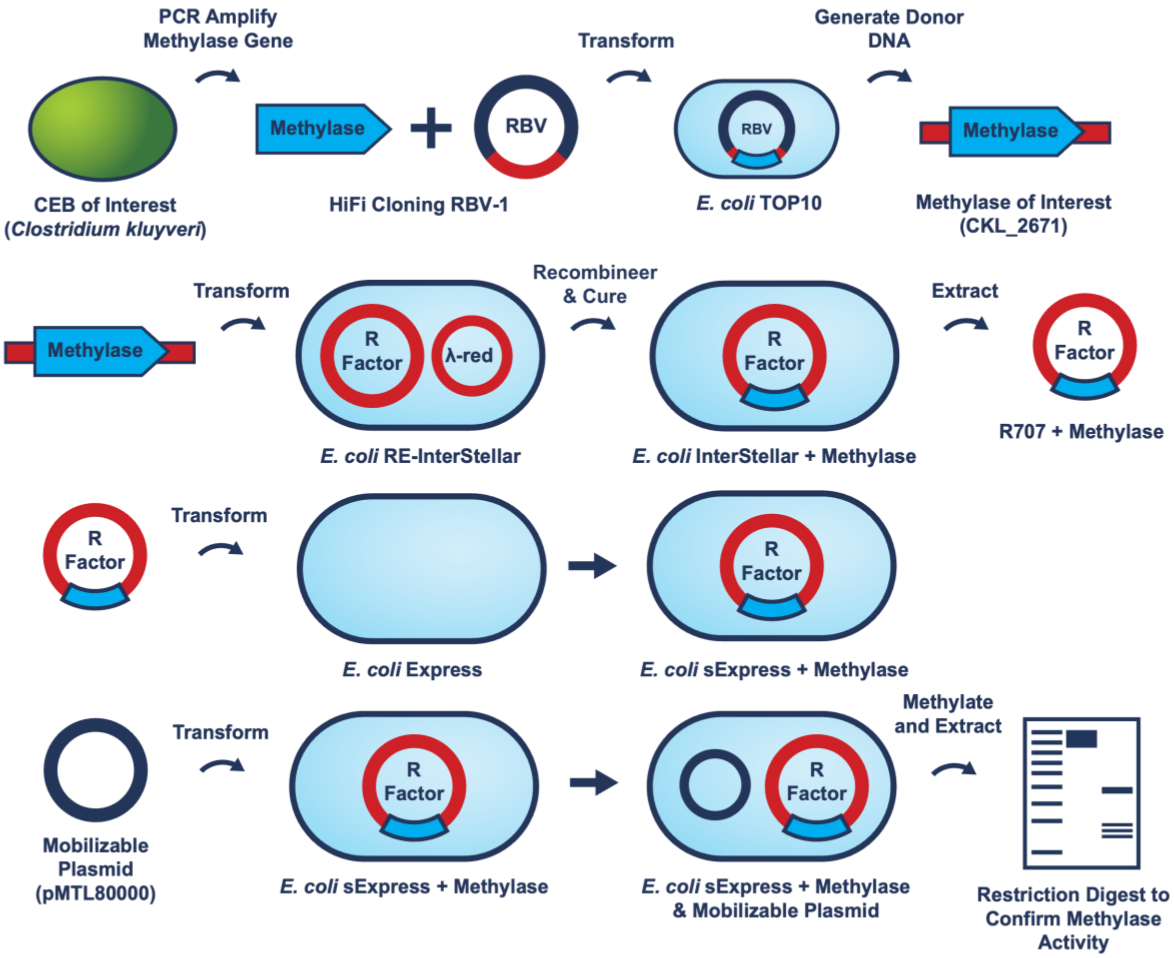


115

116 Figure S1: Schematic of the molecular cloning workflow for constructing the
117 methylating conjugal donor strain, *E. coli* sExpress CKL_2671

### Bacterial Strains, Media, and Growth Conditions

Table S2: A list of bacterial strains and their growth conditions used for this study.

| Strain | Description | Antibiotics | Temp. | Growth Media | Source |
|---|---|---|---|---|---|
| *Clostridium kluyveri* DSM555 | Type strain used for conjugation and wild-type controls | | 37°C | Hungate tubes: ATCC1120; Solid Media: Reinforced Clostridial Agar supplemented with ethanol | DSMZ |
| *Escherichia coli* NEB5a | Cloning strain | | 37°C | LB Miller broth and plates | NEB |
| *Escherichia coli* Express | Expression strain | | | | |
| *Escherichia coli* NEB5a pMTL82151 | Cloning strain carrying pMTL82151 | chloramphenicol | 37°C | LB Miller broth and plates | SBRC |
| *Escherichia coli* TOP10 RBV-1 | Cloning strain carrying RBV-1 | erythromycin | | | |
| *Escherichia coli* sExpress | Conjugal donor strain (dam+, dcm-) carrying R702 | kanamycin | | | |
| *Escherichia coli* sExpress pMTL82151 | Conjugal donor strain (dam+, dcm-) carrying R702 and pMTL82151 | kanamycin; chloramphenicol | | | |
| *Escherichia coli* RE-InterStellar | λ-red recombination strain carrying R702 and pKD46-cam; derivative of E. coli Stellar (Takara Bio) | kanamycin; chloramphenicol | 30°C | | |
| *Escherichia coli* TOP10 RBV-1-CKL_2671 | Cloning strain carrying RBV-1-CKL_2671 and used for recombineering into R702 and in vivo methylation assays | erythromycin | 37°C | LB Miller broth and plates | This work |
| *Escherichia coli* RE-InterStellar CKL_2671 | Intermediate strain carrying R702-CKL_2671 and pKD46-cam | apramycin; chloramphenicol | 30°C | | |
| *Escherichia coli* InterStellar CKL_2671 | Intermediate strain carrying R702-CKL_2671 and cured of pKD46-cam | apramycin | 37°C | | |

| | | | | | |
|---|---|---|---|---|---|
| *Escherichia coli* NEB5a pMTL82153 | Cloning strain carrying pMTL82153 and used for FAST fluorescence assays a non-fluorescent control | chloramphenicol | 37°C | LB Miller broth and plates | This work |
| *Escherichia coli* NEB5a pMTL82153-FAST | Cloning strain carrying pMTL82153-FAST and used for FAST fluorescence assays as a positive control | | | | |
| *Escherichai coli* sExpress CKL_2671 | Methylating conjugal donor strain (dam+, dcm-) carrying R702-CKL_2671 | apramycin | 37°C | LB Miller broth and plates | This work |
| *Escherichia coli* sExpress CKL_2671 pMTL82151 | Methylating conjugal donor strain (dam+, dcm-) carrying R702_CKL_2671 and pMTL82151 | apramycin; chloramphenicol | 37°C | LB Miller broth and plates | This work |
| *Escherichia coli* sExpress CKL_2671 pMTL82153 | Methylating conjugal donor strain (dam+, dcm-) carrying R702_CKL_2671 and pMTL82153 | | | | |
| *Escherichia coli* sExpress CKL_2671 pMTL82153-FAST | Methylating conjugal donor strain (dam+, dcm-) carrying R702_CKL_2671 and pMTL82153-FAST | | | | |
| *Clostridium kluyveri* DSM555 pMTL82151 | Transconjugant carrying pMTL82151 and used to validate negative selection using thiamphenicol | thiamphenicol; D-cycloserine | 37°C | Hungate tubes: ATCC1120; Solid Media: Reinforced Clostridial Agar supplemented with ethanol | This work |
| *Clostridium kluyveri* DSM555 pMTL82153 | Transconjugant carrying pMTL82153 used for FAST fluorescence assays as a non-fluorescent control | | | | |
| *Clostridium kluyveri* DSM555 pMTL82153-FAST | Transconjugant carrying pMTL82153-FAST used for FAST fluorescence assays | | | | |

**Primers**

Table S4: A list of primers used in this study.

| Name | Sequence | Template | Function |
|---|---|---|---|
| CKL_2671_F | GTTTCTCCATACCCGTTTTTTTGGGTTAAGATAAAAGGTGATTGCAATGCGTGG | *Clostridium kluyveri* DSM555 genome | Assembly of RBV-1-CKL_2671 |
| CKL_2671_R | CCGGAATCATATGACTCGAGTTTATAAAGCTATAGTTTTTATTTTTTCAATATCCTGCATTTCAGC | | |
| LHA_F | TTCTACGGCACGTTTGAAGGCG | RBV-1-CKL_2671 | Donor DNA PCR for CKL_2671 recombination into R702 |
| RHA_R | GCAGCGACTTCCACGATGGG | | |
| Int_scr_F | CCGAGCAGATCCTGCACGG | R702-CKL_2671 | Screening for R702-CKL_2671 |
| Int_scr_R | GGGAGTATGAAAAGTATGGCTGAAGCG | | |
| 82153_INS_F | GCCCCCTGCAGGATAAAAAATT | pMTL83353 | Assembly of pMTL82153 |
| 82153_INS_R | GGATTCAGAACGGCGCGCCATAAAAAT | | |
| 82153_VEC_F | ACAATTTTTTATCCTGCAGGGGGCC | pMTL82151 | |
| 82153_VEC_R | ATTTTTATGGCGCGCCGTTCTGAATCC | | |
| 82153-FAST_INS_F | TTTTAAGGAGGTGTGTTACATATGGAACACGTAGCATTTGGAAGTG | pMTL82255 | Assembly of pMTL82153-FAST |
| 82153-FAST_INS_R | TTCTTATTTTTATGCTAGCGTCATACCCTCTTAACGAAAACCCAGTAG | | |
| 82153-FAST_VEC_F | AATGGCGCTAGCATAAAAATAAGAA | pMTL82153 | |
| 82153-FAST_VEC_R | ATCATGGTCATATGTAACACACCTCCTTAAAAA | | |
| catP_scr_F | TTTGAGAGGGAACTTAGATGGTATTTGAAAAAATTG | pMTL80000 Clostridium shuttle plasmids | pMTL80000 colony PCR screening primers |
| catP_scr_R | TAAACTTAGGGTAACAAAAAACACCGTATTTCTACG | | |
| 27F | AGAGTTTGATCCTGGCTCAG | 16S rRNA gene | 16S rRNA gene amplification for strain identity verification |
| 1492R | TACCTTGTTACGACTT | | |

### Plasmids

Table S5: A list of plasmids used in this study.

| Name | Description | Selection Markers | Source |
| --- | --- | --- | --- |
| RBV-1 | Base recombineering plasmid | Erm | |
| R702 | Base conjugative transfer plasmid | Kan | SBRC |
| pKD46-cm | Temperature sensitive λ-red recombination helper plasmid | Cam | |
| RBV-1-CKL_2671 | Recombineering plasmid with CKL_2671 | Erm, Apr | |
| R702-CKL_2671 | Methylating, conjugative transfer plasmid | Kan, Apr | |
| pMTL82151 | Base pMTL82151 plasmid | Cam | This work |
| pMTL82153 | Non-fluorescent protein expression plasmid | Cam | |
| pMTL82153-FAST | Fluorescent protein expression plasmid | Cam | |

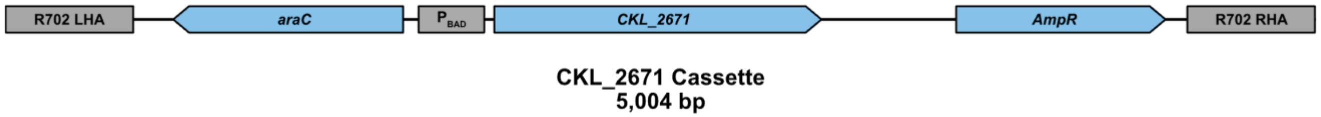


Figure S2: A schematic of the CKL_2671 Expression Cassette integrated into the R702 plasmid. R702 LHA: Left-hand homology arm (LHA) for the R702. araC: coding sequence for the $P_{BAD}$ repressor protein. $P_{BAD}$: arabinose-inducible promoter controlling CKL_2671 expression. AmpR: resistance marker for apramycin negative selection. CKL_2671: a 5mC DNA methyltransferase from the *C. kluyveri* genome, methylating **C**CGG sites at the first cytosine. R702 RHA: right-hand homology arm (RHA) for the R702.

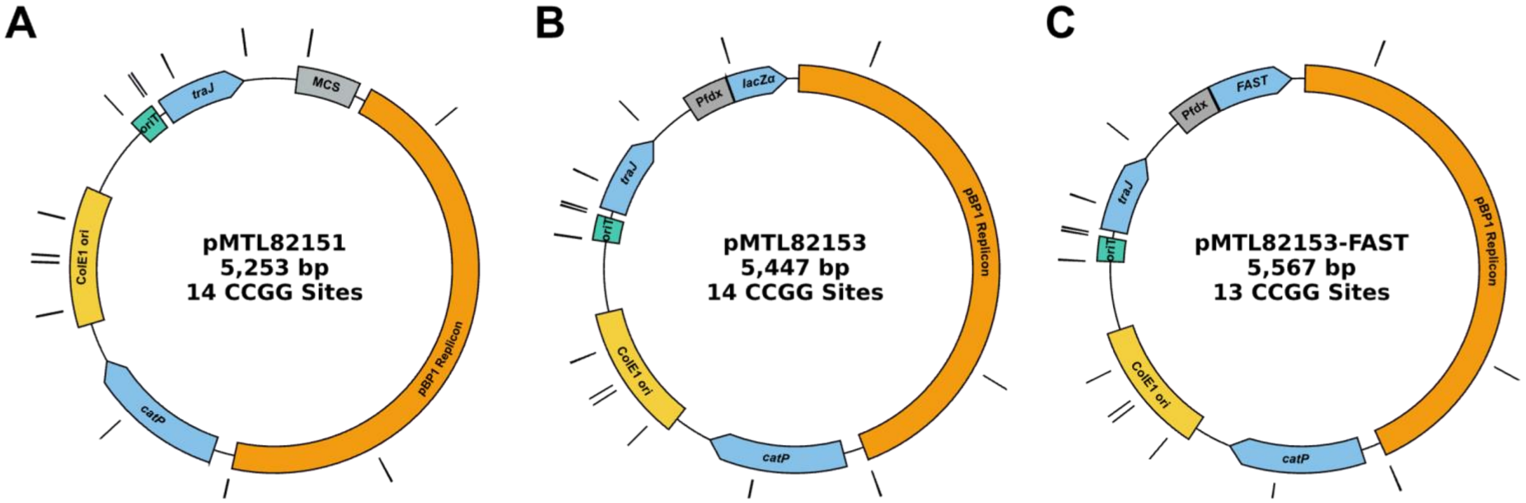


Figure S3: Plasmid maps of the pMTL80000 plasmids used in this study with annotated CCGG sites. pBP1 Replicon: Gram-positive compatible replicon from pBP1. *catP*: an antibiotic resistance marker for thiamphenicol. ColE1 ori: Gram-negative compatible origin of replication. *oriT:* an IncPβ-compatible origin of transfer. *traJ*: a coding sequence for the transcriptional activator for conjugative functions of R702. The plasmids differ at the insert region: (A) pMTL82151 contains a multiple-cloning site (MCS). (B) pMTL82153 contains an expression cassette for the lacZ α-subunit controlled by $P_{fdx}$. (C) pMTL82152-FAST contains an expression cassette for FAST controlled by $P_{fdx}$.

**Characterization of the CKL_2671 DNA Methyltransferase Activity**

Prior to constructing the methylation strain, we confirmed the *in vivo* activity and modification specificity of CKL_2671 in *E. coli*. CKL_2671 was cloned into the intermediate vector RBV-1 using HiFi Assembly. Expression of CKL_2671 in *E. coli* RBV-1 was induced using 10 mM arabinose, and plasmid DNA was extracted and subjected to MspI and HpaII digestion.

RBV-1-CKL_2671 was extracted and subjected to restriction digestion with MspI and HpaII, which both recognize the CCGG sequence targeted by the CKL_2671, methylase. These enzymes differ in their sensitivity to cytosine methylation: MspI is blocked by methylation of the external cytosine (**C**CGG), while HpaII is blocked by methylation of the internal cytosine (C**C**GG).

RBV-1-CKL_2671 was protected from cleavage by both enzymes, confirming *in vivo* activity of CKL_2671 and confirming methylation at the external cytosine (**C**CGG). The observed protection confirmed the role of CKL_2671 in the *C. kluyveri* methylome and RM system and validated it as a viable candidate for methylation strain construction.

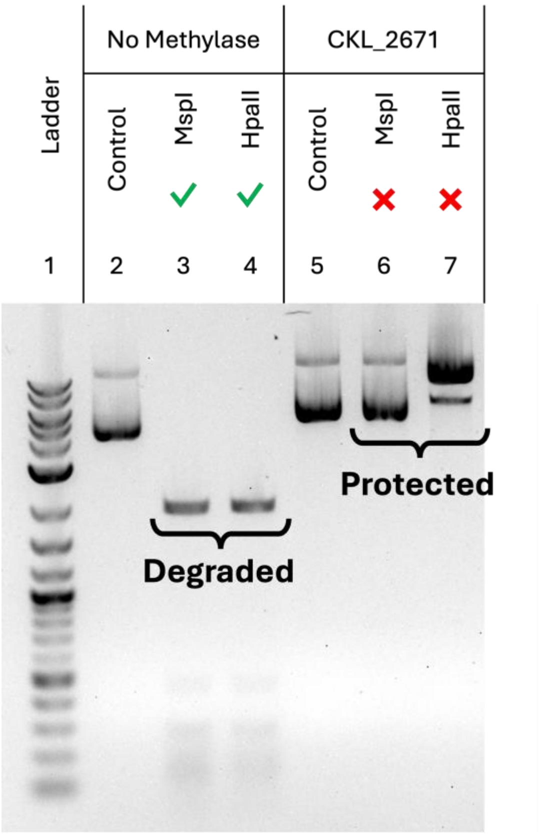


Figure S4: **1**The methylation sensitivity of the commercial restriction enzymes used in the digest. MspI and HpaII both recognize the sequence CCGG, but their activity is both blocked by methylation of the external cytosine. DNA gel electrophoresis image of plasmid digests (1% agarose). Lanes 2 and 5 are undigested controls. Lanes 3-4 and 6-7 are plasmids digested with MspI and HpaII with (RBV-1-CKL_2671) or without the methylase (RBV-1).
